# Differences in GluN2B-containing NMDA receptors result in opposite long-term plasticity and dopaminergic modulation at ipsilateral *vs*. contralateral cortico-striatal synapses

**DOI:** 10.1101/599969

**Authors:** Wei Li, Lucas Pozzo-Miller

## Abstract

Excitatory neurons in the primary motor cortex project bilaterally to the striatum. However, whether synaptic structure and function in ipsilateral and contralateral cortico-striatal pathways is identical or different remains largely unknown. Here, we describe that excitatory synapses in the contralateral pathway have higher levels of NMDA-type of glutamate receptors (NMDARs) than those in the ipsilateral pathway, although both synapses utilize the same presynaptic vesicular glutamate transporter. We also show that NMDARs containing the GluN2B subunit, but not GluN2A, contribute to this difference. The altered NMDAR subunit composition in these two pathways results in opposite synaptic plasticity: long-term depression in the ipsilateral pathway and long-term potentiation in the contralateral pathway. Furthermore, we demonstrate that activation of D1 and D2 dopamine (DA) receptors by either selective pharmacological agonists or light-induced release of endogenous DA have no effect on NMDAR-mediated neurotransmission in either pathway. However, blocking basal DAergic tone with either D1 or D2 with selective antagonists revealed that GluN2B-containing NMDARs are modulated by D1 receptors in the contralateral pathway and by D2 receptors in the ipsilateral pathway. Such distinct modulatory actions seem to be permissive rather than sufficient for the induction of long-term synaptic plasticity. Altogether, our results provide novel and unexpected evidence for the lack of bilaterality of NMDAR-mediated synaptic transmission at cortico-striatal pathways due to differences in the expression of GluN2B subunits, which results in differences in bidirectional synaptic plasticity and modulation by dopaminergic inputs.

## Introduction

In the excitatory cortico-striatal pathway, glutamatergic synapses express the characteristic complement of α-amino-3-hydroxy-5-methyl-4-isoxazolepropionic acid receptors (AMPARs) and *N*-methyl-D-aspartate receptors (NMDARs) (Shepherd, 2013; Stroebel et al., 2018). NMDARs are heteromultimers comprising two obligatory GluN1 subunits and two modulatory subunits, either GluN2 or GluN3 (Paoletti and Neyton, 2007). There are four GluN2 subunits (A-D) in the brain, but GluN2B and GluN2A predominate in the striatum (Landwehrmeyer et al., 1995; Chapman et al., 2003), where they are present in either heterodimer (GluN1/GluN2B and GluN1/GluN2A) or heterotrimeric combinations (GluN1/GluN2B/GluN2A) (Dunah and Standaert, 2003). GluN2B and GluN2A subunits are structurally and functionally distinctive, contributing unique properties to NMDAR function in basal synaptic transmission and plasticity. In addition, these two subunits are expressed at different developmental times: GluN2B is predominant in early postnatal development, whereas the levels of GluN2A progressively increase during development and ultimately exceed those of GluN2B (Monyer et al., 1994). GluN2B-containing NMDARs are preferentially targeted to extrasynaptic sites, while GluN2A-containing NMDARs are localized to the postsynaptic density (Rumbaugh and Vicini, 1999). Notably, GluN2B-containing NMDARs have lower affinity for glutamate, slower channel kinetics, and higher Ca^2+^ permeability (Erreger et al., 2005). GluN2B-containing NMDARs also have a specific binding domain for Ca^2+^/calmodulin-dependent protein kinase II (CaMKII), allowing timely activation of downstream signaling cascades that mediate long-term synaptic strengthening (Strack and Colbran, 1998). All these distinct features imparted to NMDARs by GluN2B and GluN2A subunits account for their different roles in synaptic structure and function (Shipton and Paulsen, 2013). For instance, activation of GluN2B-or GluN2A-containing NMDARs in the striatum differentially regulates GABA and glutamate release in target areas (Fantin et al., 2007), and controls glutamate and dopaminergic (DAergic) synaptic transmission (Schotanus and Chergui, 2008). Studies in the hippocampus have suggested that due to the differences in the temporal features of GluN2B-and GluN2A-mediated Ca^2+^ influx, these two subunits have a differential role in the induction of long-term potentiation (LTP) and long-term depression (LTD) (Shipton and Paulsen, 2013).

NMDAR subunits in the striatum are modulated by DAegic inputs from the substantia nigra pars compacta (SNc) (Surmeirer et al., 2007; Gerfen and Surmeier, 2011). The activation of DA receptors in the striatum mediates distinct membrane trafficking of GluN2B and GluN2A (Hallett et al., 2006). These two subunits also play a different role in DA receptor-mediated alterations of dendritic spine morphology in medium spiny neurons (MSNs) (Vastagh et al., 2012). Genetic deletion or pharmacological inhibition of GluN2A in the striatum facilitates DA receptor-mediated potentiation of NMDA responses, whereas inhibition of GluN2B prevents such potentiation (Jocoy et al., 2011). Activation of DA type 1 receptors (D1) is necessary for the induction of LTP of glutamatergic synaptic transmission (Pawlak and Kerr, 2008; Shen et al., 2008), whereas the induction of LTD requires activation of D2 receptors (Wang et al., 2006; Kreitzer and Malenka, 2007; Shen et al., 2008).

Morphological and behavioral studies have long demonstrated that cortical neurons send bilateral projections to multiple subcortical regions (Fame et al., 2011). It is generally assumed that synapses formed by ipsilateral and contralateral cortical efferents have similar postsynaptic features, and serve to synchronize inter-hemispheric activity. However, this assumption has not been directly examined. Expression of light-sensitive cation channels in cortical projection neurons allows their selective stimulation in only one hemisphere and the characterization of potential differences between ipsilateral and contralateral cortico-striatal synaptic transmission and plasticity. We uncovered the lack of bilaterality of NMDAR-mediated synaptic transmission at cortico-striatal pathways due to differences in the expression of GluN2B subunits, which results in differences in bidirectional synaptic plasticity and modulation by dopaminergic inputs.

## Materials & Methods

### Animals

*Drd1a*-tdTomato[B6.Cg-Tg(Drd1a-tdTomato)6Calak/J] (https://www.jax.org/strain/016204) and *DAT-*Cre [B6.SJL-Slc6a3tm1.1(cre)Bkmn/J] (https://www.jax.org/strain/006660) mice were purchased from The Jackson Laboratory; *Drd2*EGFP [Tg(Drd2-EGFP)S118Gsat] (http://www.informatics.jax.org/allele/MGI:3843608) mice were purchased from the Mutant Mouse Resource & Research Centers. All mice were maintained on a C57BL/6J background, and kept on a 12-h light/dark cycle with food and water *ad libitum*. Male mice (P40-70) were used for all the experiments. All animal procedures were performed in accordance and after approval by the University of Alabama at Birmingham institutional animal care and use committee.

### Stereotaxic injections

All adeno-associated viruses (AAVs) were obtained from the UNC Vector Core, and delivered via stereotaxic intracranial injections. Mice (P20-27) were anesthetized with 4% isoflurane in 100% oxygen gas; anesthesia was maintained with 1-2.5% isoflurane. Mice were placed in a stereotactic frame (David Kopf Instruments), and their body temperature was maintained with a heating pad. The scalp was shaved and then sterilized with 70% ethanol. A rostral-caudal incision was made to access the skull, a hole was drilled, and virus was delivered through a 2.5 µL syringe (Hamilton Company, Reno, NV) at a rate of 0.25 µL/min using a microsyringe pump (UMP3 UltraMicroPump, Micro4, World Precision Instruments). For optogenetic stimulation of cortical axons, AAV-CaMKIIα-ChR2 (H134R)-EYFP was unilaterally injected into the primary motor cortex (M1) (AP = +1.0 mm, ML = +1.5 mm, DV = −1.2 mm) (0.5-1 µL per site). For dual color optogenetic activation of ipsilateral and contralateral cortical inputs to the same striatal slice, AAV-Syn-Chrimson-tdTomato was injected into the ipsilateral M1, and AAV-Syn-Chronos-GFP into the contralateral M1. For optogenetic stimulation of cortical axons with high frequency theta-burst patterns, AAV-CaMKIIα-ChR2 (E123T/T159C)-EYFP (ChETA) was injected into M1. For cell-type specific conditional expression of ChR2 in DA neurons, Cre-dependent AAV-EF1α-DIO-ChR2 (H134R)-EYFP was injected into the SNc of *DAT-*Cre mice (AP = −3.0 mm, ML = ±1.3 mm, DV = −4.1 mm). Following the injections, the incision was closed with surgical glue. Topical antibiotic ointment (bacitracin zinc, neomycin sulfate, and polymyxin B sulfates; Actavis) was applied to the incision, and carprofen (5 mg/kg; Zoetis) was administered intraperitoneally. Mice were used for experiments after 3-4 weeks for the viral expression of CaMKIIα-ChR2 (H134R), Syn-Chrimson, and Syn-Chronos, and 5-6 weeks for the expression of CaMKIIα-ChETA and EF1α-DIO-ChR2.

### *Ex vivo* brain slices

Mice were deeply anesthetized with a mixture of ketamine (100 mg/kg) and xylazine (10 mg/kg), and transcardially perfused with ice-cold low Na^+^, low Ca^2+^ “cutting” artificial cerebrospinal fluid (aCSF) containing (in mM): 87 NaCl, 2.5 KCl, 0.5 CaCl_2_, 7 MgCl_2_, 1.25 NaH_2_PO_4_, 25 NaHCO_3_, 25 glucose, and 75 sucrose, bubbled with 95% O_2_/5% CO_2_. The brain was rapidly removed, and coronal (for optogenetic stimulation) or parasagittal (for electrical stimulation) slices were cut at a thickness of 300 µm with a vibrating blade microtome (VT1200S, Leica Biosystems). Coronal slices were surgically cut in the midline to separate slices ipsilateral to the AAV injection from those contralateral to the AAV injection. Slices were then transferred to normal aCSF containing (in mM): 130 NaCl, 3.5 KCl, 2 CaCl_2_, 2 MgCl_2_, 1.25 NaH_2_PO4, 24 NaHCO_3_, and 10 glucose, bubbled with 95 % O_2_ / 5 % CO_2_, at 32° C for 30 min and allowed to recover for 1 h at room temperature before recordings.

### Intracellular recordings

Individual slices were transferred to a submerged chamber mounted on a fixed-stage upright microscope (Zeiss Axioskop FS) and continuously perfused at room temperature with normal aCSF containing (in µM) 0 MgCl_2_, 50 picrotoxin (Millipore Sigma, Cat# P1675), and 10 glycine (Millipore Sigma, Cat# 8898). Slices were visualized by infrared differential interference contrast microscopy with a water-immersion 63 × objective (0.9 NA, Zeiss). D1 and D2 MSNs were identified by fluorescence excitation of tdTomato and EGFP, respectively (554 nm and 470 nm; X-Cite Turbo, Excelitas Technologies). Recordings were acquired with Axopatch-200B amplifiers (Molecular Devices), filtered at 2 kHz, and digitized at 10 kHz with an ITC-18 A/D-D/A interface (Instrutech) controlled by custom-written software in a G5 PowerMac Apple computer (TI-WorkBench, provided by Dr. Takafumi Inoue) (Inoue, 2018). Input resistance was measured with hyperpolarizing voltage pulses (50 ms, 20 mV). Cells with series resistances above 15 MΩ were discarded, and cells were also excluded if any whole-cell parameter (i.e. Cm, Ri, Rs) changed by ≥ 20% during the recordings.

Whole-cell voltage-clamp recordings were performed with unpolished pipettes (World Precision Instruments), containing (in mM): 120 Cs-gluconate, 17.5 CsCl, 10 Na-HEPES, 4 Mg-ATP, 0.4 Na-GTP, 10 Na_2_-creatine phosphate, 0.2 Na-EGTA, 5 QX-314 (290-300 mOsm, pH 7.3, final resistance: 3-4 MΩ). Total EPSCs were recorded at −50 mV in MSNs, and NMDAR EPSCs were then isolated by addition of the AMPAR antagonist 2,3-Dioxo-6-nitro-1,2,3,4-tetrahydrobenzo[*f*]quinoxaline-7-sulfonamide (NBQX, 10 µM; Tocris, Cat# 1044). NMDAR EPSCs were confirmed by addition of the NMDAR antagonist D,L-APV (20 µM; Tocris, Cat# 0106) at the end of the recordings. To evoke EPSCs in MSNs by electrical stimulation, a theta glass electrode (World Precision Instruments) was placed on the cortical layer VI close to the corpus callosum, and 100 µs-long stimuli were delivered every 20 s. To evoke EPSCs by optogenetic stimulation of ChR2 or ChETA, blue light from a fiber optic-coupled LED (470 nm, Thorlabs; 0.5-1.5 ms duration, 5-8 mW/mm^2^) was delivered to either ipsilateral or contralateral MSNs by a LED driver (LEDD1B, Thorlabs). For dual color optogenetic stimulation of Chrimson and Chronos in the same slice, red light from a fiber optic-coupled LED (625 nm, Thorlabs) was used for excitation of ipsilateral Chrimson, and blue light from a laser-LED hybrid source (470 nm, X-Cite Turbo) was used for excitation of contralateral Chronos through the 63× microscope objective. The selective GluN2B antagonist ifenprodil (3 µM; Millipore Sigma, Cat# I2892) was perfused for 10 min, followed by addition of the selective GluN2A antagonist TCN-201 (2 µM; Millipore Sigma, Cat# SML0416) for an additional 10 min. In some experiments, the selective D1 antagonist SCH-23390 (10 µM; Tocris, Cat# 0925) and the selective D2 antagonist sulpiride (10 µM; Tocris, Cat# 0895) were perfused for 10 min either before or after the GluN2B and GluN2A antagonists. The selective D1 agonist SKF-83822 (10 µM; Tocris, Cat# 2075) or the selective D2 agonist quinpirole (10 µM; Tocris, Cat# 1061) was used to activate D1 and D2 receptors.

For induction of cortico-striatal synaptic plasticity by optogenetic stimulation of cortical axons with high frequency theta-burst patterns, whole-cell current-clamp recordings of EPSPs were performed with pipettes containing (in mM): 140 K-gluconate, 5 KCl, 10 Na-HEPES, 4 Mg-ATP, 2 MgCl_2_, 0.3 Na-GTP, 10 Na_2_-creatine phosphate, 0.2 Na-EGTA (290-300 mOsm, pH 7.3, final resistance: 3-4 MΩ). Baseline EPSP amplitudes were obtained with single blue light stimuli (470 nm, Thorlabs; 0.5-1.5 ms duration, 5-8 mW/mm^2^) delivered every 30 s to either ipsilateral or contralateral MSNs at their resting membrane potential (about −80 mV). Synaptic plasticity was induced in slightly depolarized MSNs at −70mV with a theta burst pattern of blue light pulses, consisting of 10 trains of 10 bursts, each burst having 4 pulses at 100 Hz, with 200 ms between bursts, and 10 s between trains (Park et al., 2014). Pretreatment with ifenprodil or TCN-201 was used to test the requirement of GluN2B and GluN2A in ipsilateral and contralateral cortico-striatal synaptic plasticity.

### Immunofluorescence

Mice were anesthetized with a ketamine and xylazine mixture, and transcardially perfused with 4% paraformaldehyde in PBS. Brain samples were then post-fixed in 4% paraformaldehyde overnight at 4° C. Sections were cut at 60 µm using a vibrotome, permeabilized with 0.25 % Triton X-100 for 2 h, and blocked with 10 % normal serum for 1 h. Sections were incubated overnight at 4° C with anti-rabbit tdTomato (1:2000; TaKaRa, Cat# 632543, RRID:AB_2307319), anti-rabbit GFP (1:2000; Abcam, Cat# ab290, RRID:AB_2313768), anti-chicken GFP (1;2000; Abcam, Cat# ab13970, RRID:AB_300798), anti-rabbit mCherry (1:2000; Abcam, ab167453, RRID:AB_2571870), anti-guinea pig VGLUT1 (1:500; Millipore Sigma, Cat# AB5905, RRID:AB_2301751), anti-rabbit VGLUT2 (1:500; Millipore Sigma, Cat# V-2514, RRID:AB_477611), anti-mouse tyrosine hydroxylase (TH; 1:1000; Millipore Sigma, Cat# MAB318, RRID:AB_2313764), or anti-rabbit NeuN (1:1000; Millipore Sigma, Cat# ABN78, RRID:AB_10807945) primary antibodies. After the primary antibodies, sections were rinsed with PBS three times each 10 min and incubated for 2 h at room temperature with corresponding secondary antibodies tagged with Alexa Fluor-488, −594, or −647 (Jackson ImmunoResearch). Sections were coverslipped with Vectashield mounting media (Vector Laboratories). All images were acquired using 10× (0.3 NA), 20× (0.8 NA), or 63× (1.4 NA) objectives in an LSM-800 confocal microscope (Zeiss).

### Statistical analyses

All data were analyzed using Prism (GraphPad). Comparisons between groups were analyzed by two-tailed unpaired student’s *t* test, or non-parametric Mann-Whitney test. Two-way ANOVA repeated measures were used for comparing effect of treatments for some data as stated. All data are shown as mean ± standard error, with *n* and specific statistical test as described in the figures or text. Differences were considered statistically significant at \**p* < 0.05, ** *p* < 0.01, and *** *p* < 0.0001. Statistical Power was calculated with G*Power (Faul et al., 2009).

## Results

To selectively recruit ipsilateral or contralateral cortical efferents onto striatal MSNs, we injected AAVs into one hemisphere of the M1 for the expression of CaMKIIα-driven channelrhodopsin-2 (ChR2) in excitatory cortical neurons (Fig. 1*A*, asterisk). Immunostaining for the marker EYFP demonstrates that, in addition to a dense innervation in the ipsilateral striatum, labeled M1 neurons also send a decussating callosal projection to the contralateral striatum. In the striatum, MSNs receives diverse excitatory inputs that express either vesicular glutamate transporter 1 (VGLUT1) or 2 (VGLUT2) in their presynaptic terminals (Fremeau et al., 2001; Herzog et al., 2001). Double immunostaining for either VGLUT1 or VGLUT2 and EYFP showed that VGLUT1, but not VGLUT2, co-localized with EYFP in the ipsilateral and contralateral pathways (Fig. 1*B*), indicating that both cortical pathways use the same presynaptic vesicular glutamate transporter.

**Figure 1.**
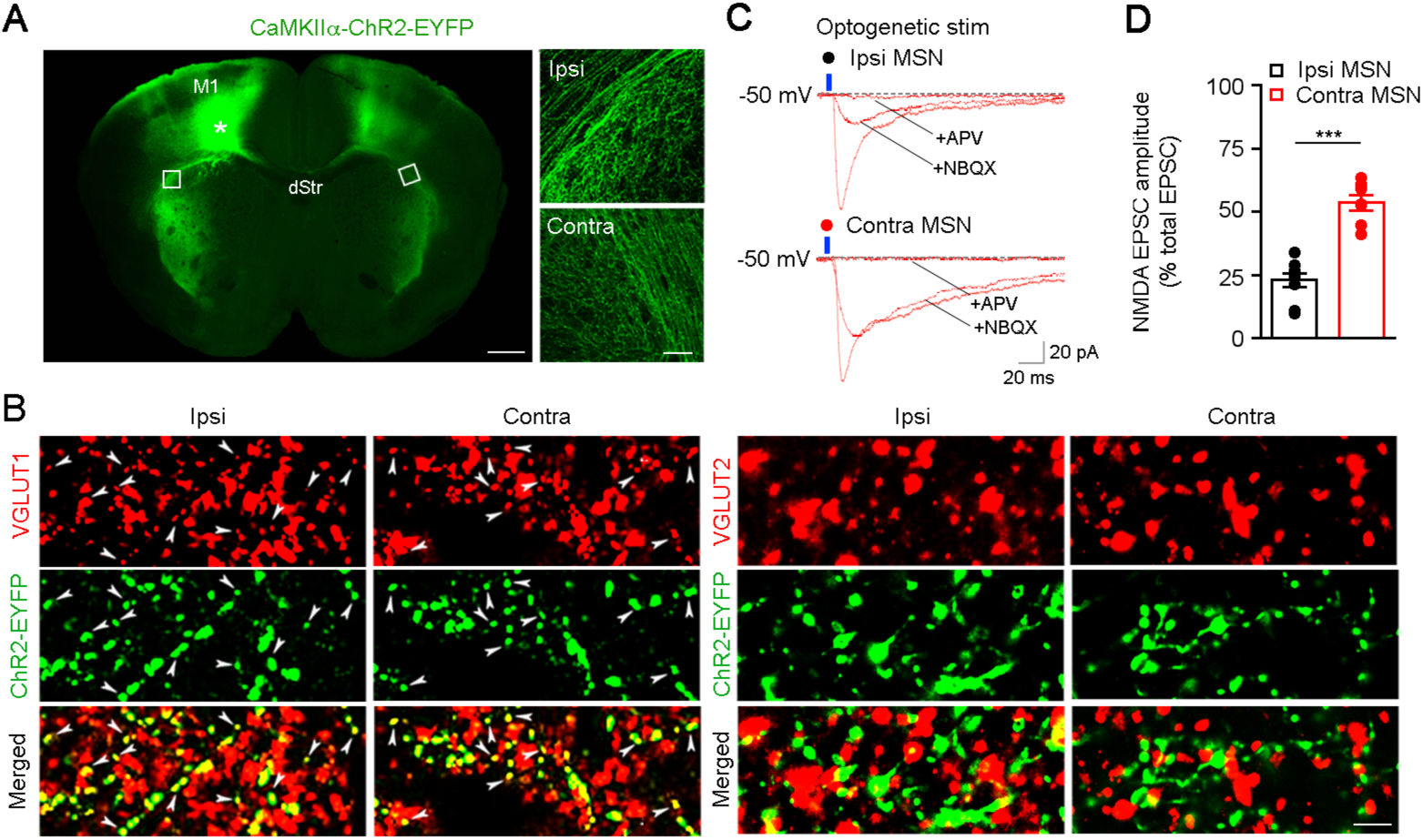
Morphological and functional features of ipsilateral and contralateral cortico-striatal pathways. ***A***, EYFP immunostaining shows the bilateral projection of M1 cortical neurons in the striatum. Asterisk shows the injection site of AAV-CaMKIIα-ChR2-EYFP in M1; white boxes in the ipsilateral and contralateral dorsolateral striatum are enlarged at right. Scale bars: 500 µm (*left*), 50 µm (*right*). ***B***, Double VGLUT1 and EYFP immunostaining (*left*) shows co-localized puncta (arrowheads) in the striatum of both hemispheres, ipsilateral and contralateral to the AAV-CaMKIIα-ChR2-EYFP injection site in M1. Double VGLUT2 and EYFP immunostaining (*right*) shows a lack of co-localization. Scale bar: 5 µm. ***C***, Representative EPSCs (from ***D***) evoked by blue light (470 nm) in ipsilateral (*top*) and contralateral (*bottom*) MSNs in striatal slices from mice expressing CaMKIIα-ChR2-EYFP in M1. Traces are from baseline, 10 min after NBQX, and 10 min after D,L-APV. ***D***, Average of the NMDAR component of the blue light-evoked EPSC in ipsilateral and contralateral MSNs (n = 9, Ipsi MSN; n = 7, Contra MSN; \*\*\**p* < 0.0001, unpaired Student’s *t* test).

To pharmacologically isolate excitatory postsynaptic currents (EPSCs) mediated by NMDARs in MSNs, blue light (470 nm) was delivered to coronal striatal slices perfused with Mg^2+^-free aCSF containing the NMDAR modulator glycine, the AMPAR antagonist NBQX, and the GABA_A_R antagonist picrotoxin (Fig. 1*C*). NMDAR-mediated EPSCs were confirmed by their complete blockade by the NMDAR antagonist D,L-APV. We compared the amplitude of NMDAR-mediated EPSCs in ipsilateral MSNs vs. contralateral MSNs. Unexpectedly, the fraction of the ESPC amplitude mediated by NMDARs (normalized to the total EPSC amplitude) is significantly larger in contralateral (53.5±3.2%) than in ipsilateral (22.9±2.7%) MSNs (Fig. 1*D*).

Because (1) GluN2B and GluN2A subunits are the major components of NMDARs in the striatum (Landwehrmeyer et al., 1995; Chapman et al., 2003), and (2) the striatum contains two distinct types of neurons, D1-expressing MSNs and D2-expressing MSNs (Gerfen and Surmeier, 2011), we examined the properties conferred to NMDARs by these two subunits at cortico-striatal synapses in *ex vivo* slices from double transgenic mice expressing tdTomato in D1 neurons and EGFP in D2 neurons (*Drd1a*-tdTomato::*Drd2*-EGFP) (Fig. 2*A*).

**Figure 2.**
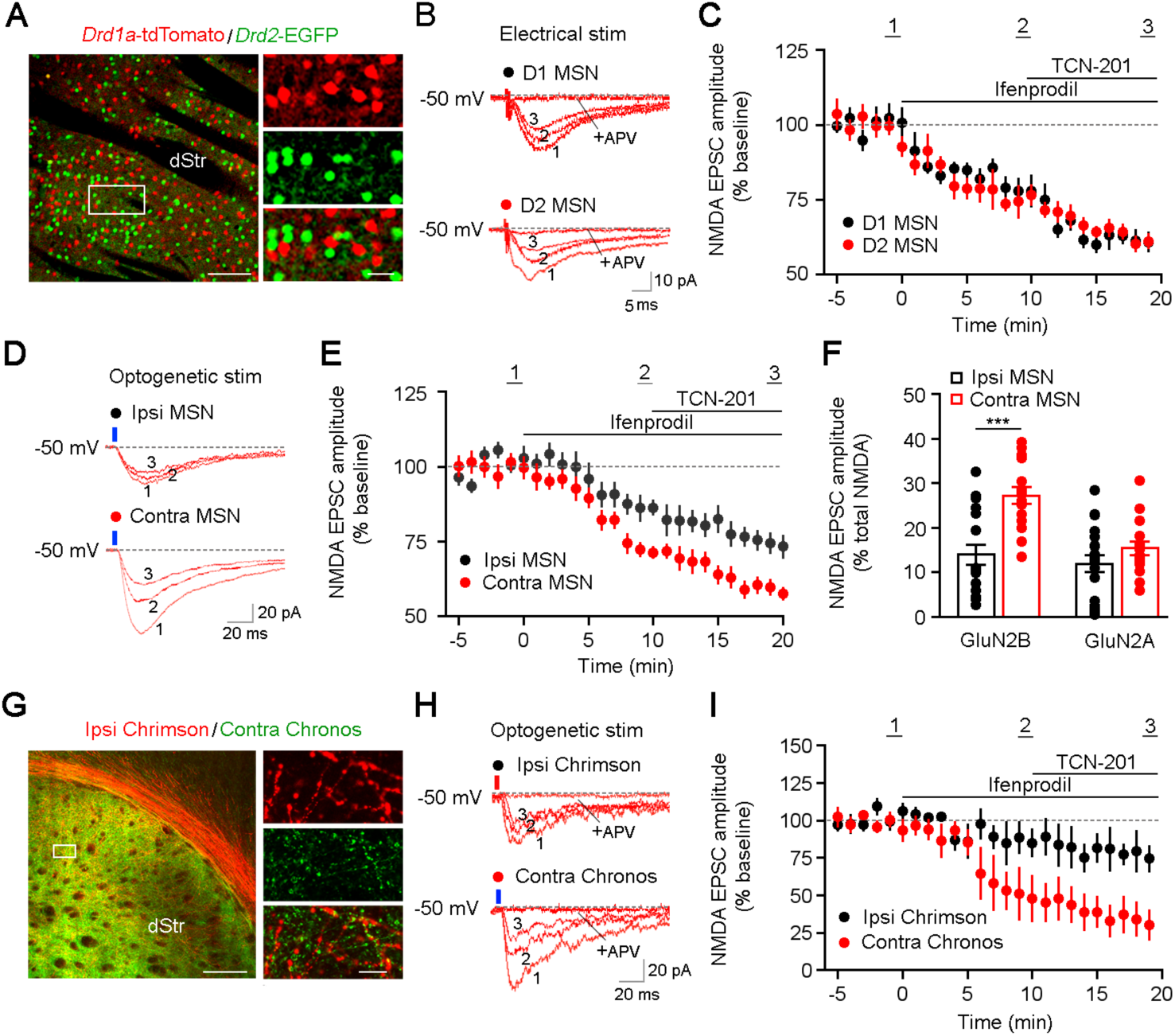
Differences in subunit composition of synaptic NMDARs between the ipsilateral and contralateral cortico-striatal pathways. ***A***, Double tdTomato and EGFP immunostaining shows segregation of D1 and D2 MSNs in the striatum of *Drd1a*-tdTomato::*Drd2*-EGFP double transgenic mice. ***B***, Representative NMDAR EPSCs (from ***C***) in D1 (*top*) and D2 MSNs (*bottom*) evoked by electrical stimulation of the cortico-striatal pathway. Traces are from baseline (1), 10 min after ifenprodil (2), 10 min after TCN-201 (3), and 10 min after D,L-APV. ***C***, Average effects of ifenprodil, TCN-201, and D,L-APV on NMDAR EPSC amplitudes (n = 7, D1 MSN; n = 6, D2 MSN; *p* = 0.56, two-way ANOVA). ***D***, Representative EPSCs (from ***E***) evoked by blue light in ipsilateral (*top*) and contralateral (*bottom*) MSNs. Traces are from baseline (1), 10 min after ifenprodil (2), and 10 min after TCN-201 (3). ***E***, Average effects of ifenprodil and TCN-201 on blue light-evoked NMDAR EPSCs (n = 19, Ipsi MSN; n = 16, Contra MSN; *p* = 0.0013, two-way ANOVA). ***F***, Average percentage of the ifenprodil-and TCN-201-sensitive components of NMDAR-mediated EPSCs. The ifenprodil-sensitive component of the NMDAR EPSCs was calculated by subtracting EPSCs after 10 min in ifenprodil from baseline EPSCs; the TCN-201-sensitive component of the NMDAR EPSCs was calculated by subtracting EPSCs after 10 min in TCN-201 from EPSCs in the presence of ifenprodil (\*\*\**p* < 0.0001, unpaired student’s *t* test). ***G***, tdTomato and GFP immunostaining shows ipsilateral Chrimson expression and contralateral Chronos expression. Scale bars: 100 µm (*left*), 10 µm (*right*). ***H***, Representative EPSCs (from ***I***) evoked by either Chrimson-activating red light (625 nm, *top*) or Chronos-activating blue light (470 nm, *bottom*). Traces are from baseline (1), 10 min after ifenprodil (2), and 10 min after TCN-201 (3). ***l***, Average effects of ifenprodil and TCN-201 on red light-and blue light-evoked NMDAR EPSCs (n = 19, Ipsi MSN; n = 16, Contra MSN; *p* = 0.028, two-way ANOVA).

We first recorded pharmacologically isolated NMDAR-mediated EPSCs in MSNs evoked by electrical stimulation of M1 layer VI close to the corpus callosum. The amplitude of NMDAR-mediated EPSCs was reduced by the selective GluN2B antagonist ifenprodil, and further reduced by addition of the GluN2A antagonist TCN-201 (Fig. 2*B*). The fractional reductions by ifenprodil and TCN-201 were comparable in D1 MSNs and D2 MSNs (Fig. 2*C*). To more unequivocally demonstrate if the presence of GluN2B and/or GluN2A subunits is responsible for the difference in the amplitude of NMDAR-mediated EPSCs in the ipsilateral and contralateral pathways, we used *ex vivo* brain slices from mice expressing ChR2 in M1 of one brain hemisphere. The reductions of NMDAR-mediated EPSCs by sequential application of ifenprodil and TCN-201 were significantly larger in MSNs contralateral to the ChR2-expressing M1 than in those MSNs ipsilateral to the labeled M1 (Fig. 2*D,E*). In addition, the fraction of the EPSC mediated by GluN2B-contaning NMDARs was significantly larger in contralateral MSNs (*p* < 0.0001), while that of the EPSC mediated by GluN2A-contaning NMDARs was not (*p* = 0.18) (Fig. 2*F*). To directly compare ipsilateral and contralateral cortical inputs on MSNs of the same striatal slice, we expressed the blue light-activated opsin Chronos in M1 of one hemisphere and the red light-activated opsin Chrimson in the opposite M1, and recorded NMDAR EPSCs in MSNs in one side of the striatal slice (Fig. 2*G*). Consistently, the effects of sequential application of ifenprodil and TCN-201 on the amplitude of NMDAR-mediated EPSCs were larger for blue light-stimulated contralateral efferents than for red light-stimulated ipsilateral efferents, and that reduction is due to NMDARs containing GluN2B subunits (Fig. 2*H*,*l*).

We next tested whether different levels of GluN2B-and GluN2A-contaning NMDARs at ipsilateral and contralateral cortico-striatal synapses could result in distinct synaptic plasticity properties. To selectively activate axons from M1 pyramidal neurons from only one hemisphere with theta-burst stimulation (TBS) light patterns, we use ChETA because it follows high frequency light pulse trains with less inactivation than ChR2 (Xie et al., 2013). TBS pattern of blue light pulses reliably induced temporally summating subthreshold excitatory postsynaptic potentials (EPSPs) in MSNs (Fig. 3*A*). As expected, blue-light TBS of cortical inputs to contralateral MSNs resulted in LTP of EPSP amplitudes (133.7±3.6% of baseline, 25 min after TBS) (Fig. 3*C*), which was blocked by either ifenprodil or TCN-201. Surprisingly, blue-light TBS of cortical inputs to ipsilateral MSNs resulted in LTD of EPSP amplitudes (54.8±5.5% of baseline, 25 min after TBS) (Fig. 3*B*), which was also blocked by either ifenprodil or TCN-201. These data demonstrate that GluN2B-and GluN2A-contaning NMDARs are both required for LTP and LTD in ipsilateral and contralateral cortico-striatal pathways, but the differential content of GluN2B subunits of synaptic NMDARs determined the direction of long-term synaptic plasticity.

**Figure 3.**
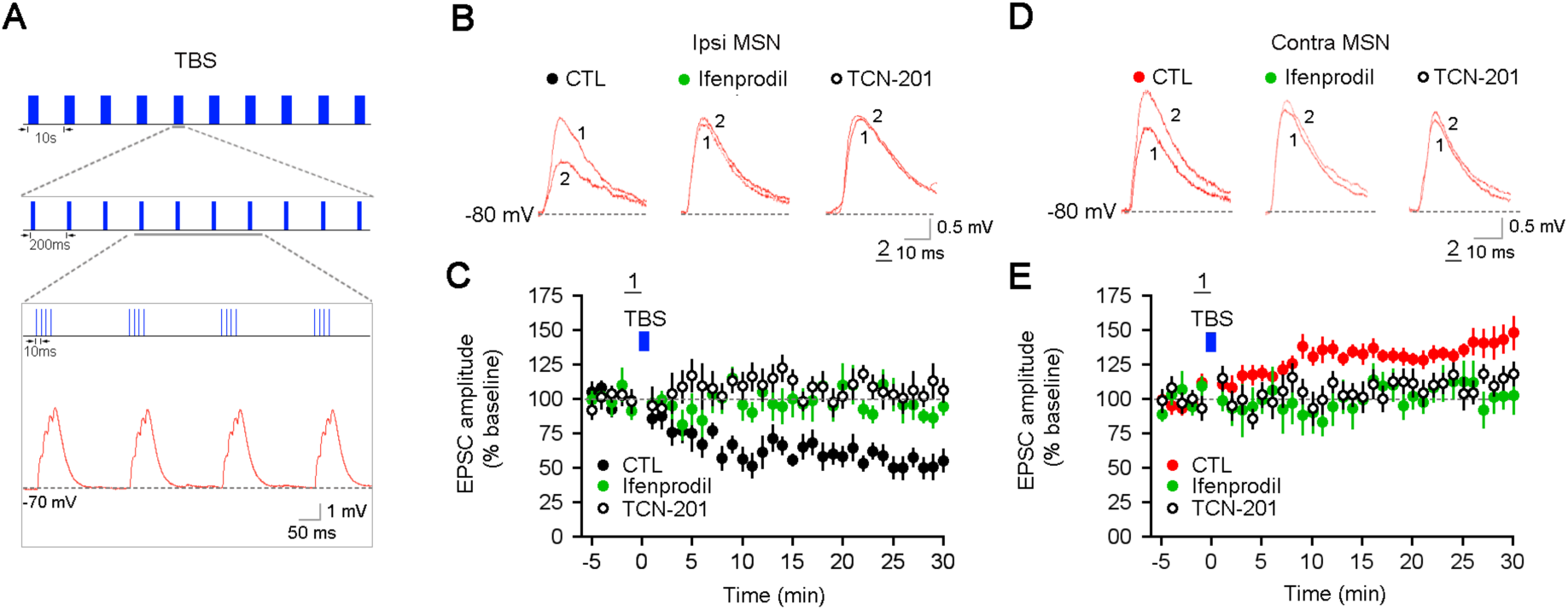
The role of GluN2A-and GluN2B-containing NMDARs in synaptic plasticity at the ipsilateral and contralateral cortico-striatal pathways. ***A***, Theta-burst pattern of blue light (470 nm) pulses to activate ChETA in axons of M1 pyramidal neurons in a striatal slice (*top*) evokes temporally summating subthreshold EPSPs in a MSN (*bottom*). ***B***, Representative EPSPs (from ***C***) evoked by a single blue light pulse before (1) and 25 min after TBS of blue light pulses (2) in control or slices treated with either ifenprodil or TCN-201. ***C***, TBS of blue light (bar) induced LTD of EPSPs in ipsilateral MSNs, which was blocked by either ifenprodil or TCN-201 (n = 7, CTL; n = 8, ifenprodil; n = 9, TCN-201; *p* < 0.0001 CTL vs. ifenprodil and TCN-201, two-way ANOVA). ***D***, Representative EPSPs (from ***E***) evoked by a single blue light pulse before (1) and 25 min after TBS of blue light pulses (2) in control or slices treated with either ifenprodil or TCN-201. ***E***, TBS of blue light (bar) induced LTP of EPSPs in contralateral MSNs, which was blocked by either ifenprodil or TCN-201 (n = 7, CTL; n = 7, ifenprodil; n = 10, TCN-201; *p* = 0.0023 ifenprodil vs. CTL; *p* = 0.0049 TCN-201 vs. CTL; two-way ANOVA).

Because glutamatergic synapses between M1 pyramidal neurons and striatal MSNs are modulated by DAergic inputs from the SNc (Gerfen and Surmeier, 2011), we next characterized the modulation of blue light-evoked NMDAR-mediated EPSCs in MSNs by using selective D1 and D2 receptor agonists. Neither the D1 agonist SKF-83822 (Fig. 4*A,B*) nor the D2 agonist quinpirole (Fig. 4*C,D*) had any effect on the amplitude of NMDAR-mediated EPSCs evoked in ipsilateral or contralateral MSNs. To directly test the actions of endogenously released DA on these responses, we injected AAVs expressing Cre-dependent ChR2 in the SNc of *DAT-*Cre mice, which resulted in its selective expression in DAergic neurons. Immunostaining for the marker EYFP in striatal sections showed abundant afferent DAergic fibers originating in the SNc (Fig. 4*E*). Compared to baseline conditions without blue-light stimulation (Fig. 4*F*), the amplitude of NMDAR-mediated EPSCs evoked by electrical stimulation of cortical afferent was not affected by either short blue light trains (three 5 ms pulses delivered every 15 ms; Fig. 4*G*), or a long train of light stimuli (a single short burst delivered 25 times every 1 s; Fig. 4*H*) designed to model the phasic firing mode of SNc DAergic neurons.

**Figure 4.**
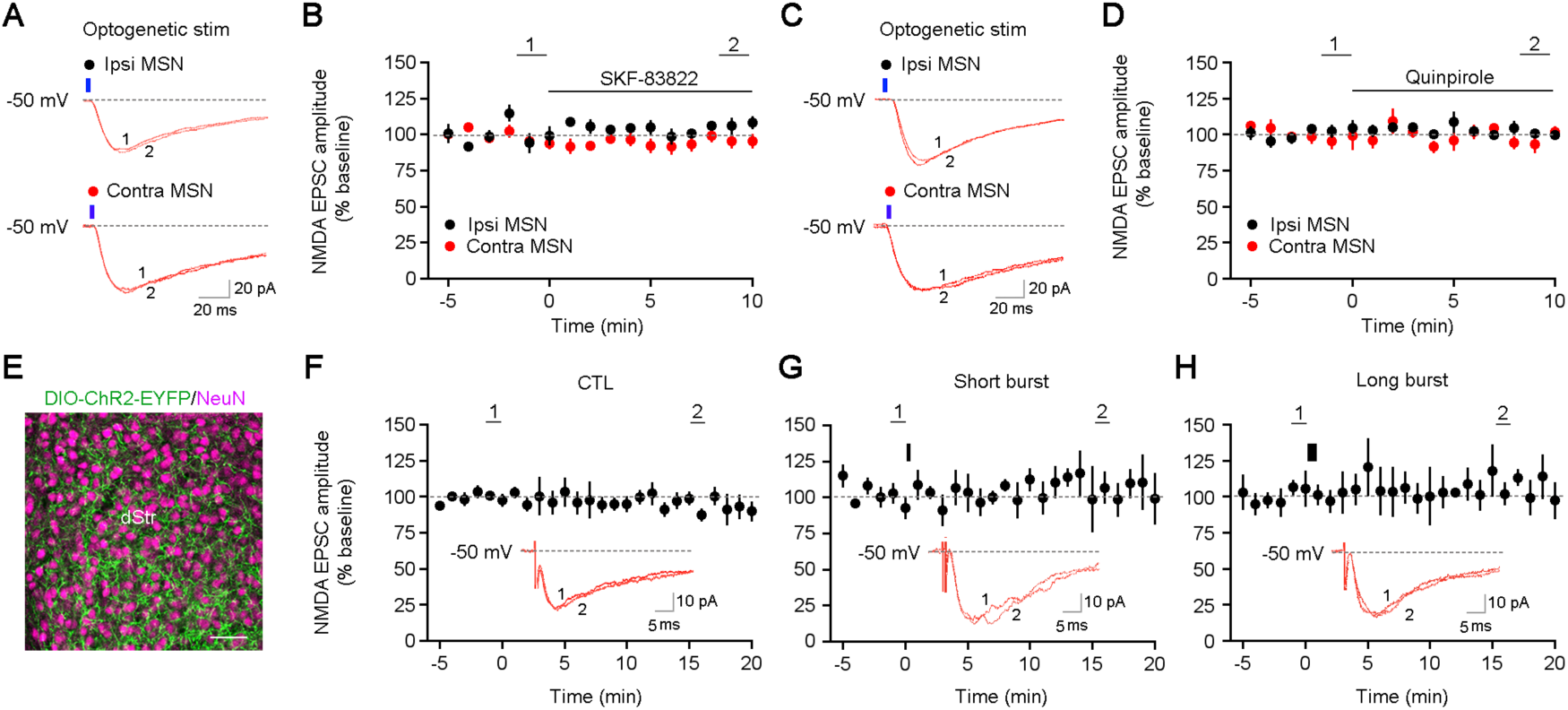
Activation of D1 and D2 receptors by either selective agonists or optogenetic-induced endogenous DA release has no effect on NMDAR-mediated EPSCs at ipsilateral and contralateral pathways. ***A***, Representative EPSCs (from ***B***) evoked by blue light (470 nm) in ipsilateral (*top*) and contralateral (*bottom*) MSNs. Traces are taken at baseline (1), and 10 min after SKF-83822 (2). ***B***, Average effect of SKF-83822 on blue light evoked NMDAR EPSCs (n = 4, Ipsi MSN; n = 7, Contra MSN; *p* = 0.06, two-way ANOVA). ***C***, Representative EPSCs (from ***D***) evoked by blue light in ipsilateral (*top*) and contralateral (*bottom*) MSNs. Traces are taken at baseline (1), and 10 min after quinpirole (2). ***D***, Average effect of quinpirole on blue light evoked NMDAR EPSCs (n = 4, Ipsi MSN; n = 6, Contra MSN; *p* = 0.19, two-way ANOVA). ***E***, AAV-EF1α-DIO-ChR2(H134R)-EYFP was injected into the SNc of *DAT-*Cre (B6.SJL-Slc6a3tm1.1(cre)Bkmn/J) mice for the expression of ChR2 in DA neurons. Dual EYFP and NeuN immunostaining in the dorsal striatum (dStr) shows afferent fibers (green) from the SNc. ***F-H***, Average amplitudes of electrically evoked NMDAR EPSCs in MSNs without (***F***; CTL, n=5), and after endogenous DA release by either a short burst (***G***; narrow black bar, three 5 ms pulses delivered every 15 ms; n = 5), or a long burst of blue light stimuli (***H***; wide black bar, a single short burst delivered 25 times every 1 s; n = 4). Insets show representative electrically evoked NMDAR EPSCs; traces are taken at baseline (1), and 10 min (2) later without light stimulation, or 10 min after a short or a long burst of blue light stimuli to release DA from afferent fibers in the striatum.

We finally tested whether application of selective D1 and D2 antagonists revealed a tonic activation of DA receptors modulating cortico-striatal NMDAR-mediated synaptic transmission. Indeed, the selective D1 antagonist SCH-23390 reduced the amplitude of NMDAR-mediated EPSCs evoked by blue light in ipsilateral MSNs (Fig. 5*A,C*), without changing the proportion of the EPSC mediated by GluN2B-and GluN2A-containing NMDARs, as determined by sequential application of ifenprodil and TCN-201. Similar results were observed when SCH-23390 was applied after ifenprodil and TCN-201 had already reduced the amplitude of NMDAR-mediated EPSCs (Fig. 5*B,C*). On the other hand, the D2 antagonist sulpiride increased the amplitude of NMDAR-mediated EPSCs in ipsilateral MSNs (117.6±4.6% Fig. 5*D,F*), which was followed by a significantly larger reduction in NMDAR-mediated EPSCs by ifenprodil (32.8±4.5% compare with 14.0±2.3% in Fig. 1*F, p* < 0.0001, unpaired Student’s *t* test). However, sulpiride had no effect on EPSCs when applied after ifenprodil and TCN-201 had already reduced the amplitude of NMDAR-mediated EPSCs (Fig. 5*E,F*). In contralateral MSNs, blocking D1s with SCH-23390 did not affect blue light-evoked NMDAR-mediated EPSCs (Fig. 5*A,C*), but subsequent application of ifenprodil resulted in a significantly smaller reduction of NMDAR-mediated EPSC amplitudes (18.2±4.1% compare with 27.3±1.9% in Fig. 1*F, p* = 0.031, unpaired Student’s *t* test). However, SCH-23390 did reduce the amplitude of NMDAR-mediated EPSCs when applied after they had already been reduced by ifenprodil and TCN-201 (Fig. 5*B,C*). On the other hand, blocking D2Rs with sulpiride did not affect the amplitude of NMDAR-mediated EPSCs either before or after inhibition of GluN2B-containing NMDARs with ifenprodil and GluN2A-containing NMDARs with TCN-201 (Fig. 5*D-F*).

**Figure 5.**
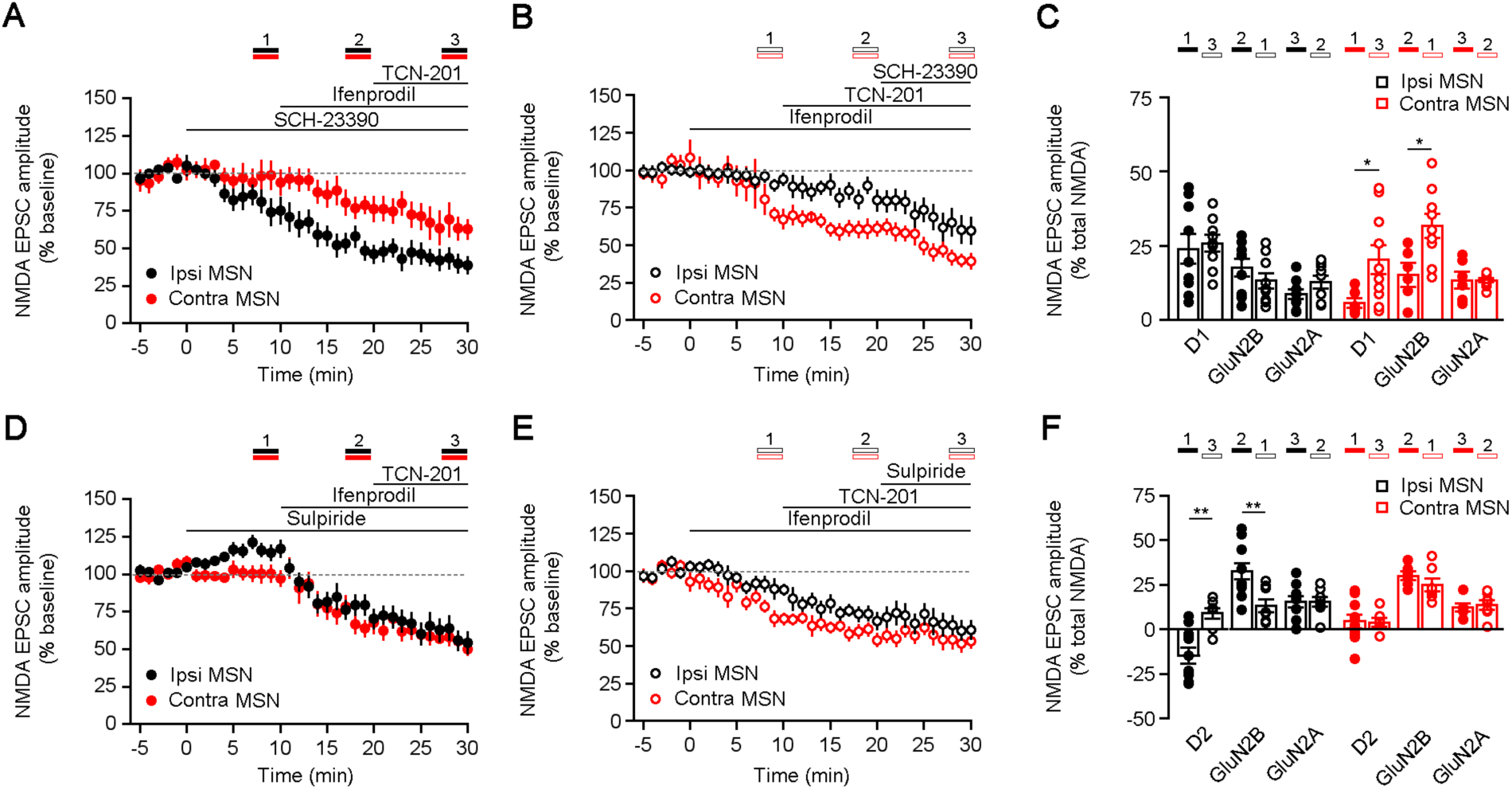
Tonic D1 and D2 activation differentially modulate GluN2B-containing NMDARs in the ipsilateral and contralateral cortico-striatal pathways. ***A***, Average effects of SCH-23390 on blue light-evoked NMDAR EPSCs in ipsilateral and contralateral MSNs, followed by ifenprodil and TCN-201 (n = 9, Ipsi MSN; n = 6, Contra MSN; *p* = 0.027, two-way ANOVA). ***B***, Average effects of SCH-23390 when applied after ifenprodil and TCN-201 had reduced blue light-evoked NMDAR EPSCs (n = 10, Ipsi MSN; n = 10, Contra MSN; *p* = 0.027, two-way ANOVA). ***C***, Average percentage of SCH-23390-, ifenprodil-, and TCN-201-sensitive components of the NMDAR-mediated EPSC. First, the SCH-23390-sensitive component of the NMDAR EPSCs was calculated by subtracting EPSCs after 10 min in SCH-23390 from baseline EPSCs. Next, the ifenprodil-sensitive component of the NMDAR EPSCs was calculated by subtracting EPSCs after 10 min in ifenprodil from EPSCs in the presence of SCH-23390. Finally, the TCN-201-sensitive component of the NMDAR EPSCs was calculated by subtracting EPSCs after 10 min in TCN-201 from EPSCs in the presence of SCH-23390 and ifenprodil (\**p* < 0.05, D1, Mann Whitney test; GluN2B, unpaired Student’s *t* test). The similar calculation was performed for the reserve treatments. ***D***, Average effects of sulpiride on blue light-evoked NMDAR EPSCs in ipsilateral and contralateral MSNs, followed by ifenprodil and TCN-201 (n = 17, Ipsi MSN; n = 10, Contra MSN; *p* = 0.32, two-way ANOVA). ***E***, Average effects of sulpiride when applied after ifenprodil and TCN-201 had reduced blue light-evoked NMDAR EPSCs (n = 7, Ipsi MSN; n = 8, Contra MSN; *p* = 0.036, two-way ANOVA). ***F***, Average percentage of SCH-23390-, ifenprodil-, and TCN-201-sensitive components of the NMDAR-mediated EPSCs; each component was calculated as described in ***C*** (\*\**p* < 0.01, D2, Mann Whitney test; GluN2B, unpaired student’s *t* test).

## Discussion

In this study, we provide the first evidence that the glutamatergic synapses in the contralateral cortico-striatal pathway contain higher levels of GluN2B-containing NMDARs than those in the ipsilateral pathway. We also demonstrate that such distinct NMDAR composition results in LTP at contralateral cortico-striatal synapses, but LTD at ipsilateral cortico-striatal synapses following the same pattern of optogenetic stimulation of ChETA-expressing axons of M1 pyramidal neurons. Finally, we show that tonic activation of D1 receptors regulate GluN2B-containing NMDARs at contralateral cortico-striatal synapses, while they are subject to D2 modulation at ipsilateral cortico-striatal synapses. The amplitude of intracellular Ca^2+^ levels triggered by influx through NMDARs dictates the induction of LTP and LTD, with higher Ca^2+^ levels promoting LTP (Shouval et al., 2002). Consistently, we observed that the levels of NMDARs in the cortico-striatal pathways correlated with the direction of synaptic plasticity, with LTP in the contralateral pathway that express higher NMDAR levels, and LTD in the ipsilateral pathway that express lower NMDAR levels. In addition, we found that such distinct consequences are mainly due to different expression levels of GluN2B but not GluN2A. Studies in the hippocampus have suggested that the differential contribution of GluN2A-or GluN2B-containing NMDARs to LTP and LTD varies, depending on the stimulus paradigm, developmental stage, and the specificity and concentration of the pharmacological antagonists used (Shipton and Paulsen, 2013). An early study in the hippocampus showed that antagonism of GluN2A-containing NMDARs prevented the induction of high frequency stimulation-and pairing-induced LTP, but antagonism of GluN2B-containing NMDARs prevented the induction of LTD by low-frequency afferent stimulation (Liu et al., 2004). In young rats, both LTP and LTD were reduced by antagonism of GluN2A-containing NMDARs, but only LTP was decreased by antagonism of GluN2B-containing NMDARs (Bartlett et al., 2007). Moreover, spike-timing-dependent LTP, but not high-frequency stimulation-induced LTP, was prevented by an antagonist of GluN2B-containing NMDARs (Zhang et al., 2008); however, higher concentration of this antagonist also impaired TBS-induced LTP (Volianskis et al., 2013). Genetic studies show that enhancing GluN2B expression in the hippocampus led to an increase in LTP and not LTD (Tang et al., 1999; Wang et al., 2009), while GluN2B knockout or RNAi-mediated GluN2B knockdown resulted in an impairment of LTP and LTD (Kutsuwada et al., 1996; Akashi et al., 2009; Foster et al., 2010). In addition, genetic deletion of GluN2A subunits resulted in a decreased in LTP and LTD amplitude in the dentate gyrus (Kannangara et al., 2015). In the striatum, GluN2B and GluN2A subunits are thought to differentially shape the time window for the induction of LTP and LTD during spike-timing-dependent plasticity (Evans et al., 2012). Our results demonstrate that higher levels of GluN2B-containing NMDAR at contralateral cortico-MSN synapses are associated with LTP, whereas lower levels of these receptors at ipsilateral cortico-MSN synapses are associated with LTD. Our findings also suggest even though the levels of GluN2B-containing NMDAR define the polarity of plasticity, NMDAR containing both subunits are still needed for LTP and LTD, as antagonists of GluN2B-or GluN2A-containing NMDARs prevent long-term changes in EPSP amplitude after afferent stimulation. In addition to GluN1/GluN2A and GluN1/GluN2B, there exists GluN1/GluN2A/GluN2B in the striatum (Li et al., 2004). The role of NMDAR triheteromers needs to be further explored, because the pharmacological properties of the triheteromers substantially distinguish from those of the diheteromers (Stroebel et al., 2018).

Induction of long-term synaptic plasticity in striatal slices has been challenging (Lovinger, 2010). The high-frequency stimulation pattern used to induce LTP often results in the induction of LTD, and vice versa. This difficulty has been ascribed to an excessive blockade of NMDARs by extracellular Mg^2+^, and the alteration of intracellular signaling during whole-cell recordings, as experiments in extracellular solution with low Mg^2+^ and the use of extracellular field recordings improve the success rate of LTP and LTD induction. Our results provide an additional explanation for the inconsistent observations of plasticity in striatal MSNs. The use of electrical electrodes inevitably results in the stimulation of axons coming from the ipsilateral and the contralateral primary motor cortex, which confounds the results due to the different levels of GluN2B-containing NMDARs in each pathway. In this condition the direction of plasticity is uncertain, in contrast to clear optogenetically-induced plasticity,

Similar to the distinct role of the different NMDAR composition in synaptic plasticity, D1 and D2 receptors have been differentially implicated in LTP and LTD of cortico-striatal glutamatergic transmission (Surmeirer et al., 2007; Gerfen and Surmeier, 2011). Consistently, we observed that D1 receptors modulate GluN2B-containing NMDARs in the contralateral pathway (which expresses LTP), while D2 receptors modulate GluN2B-containing NMDARs in the ipsilateral pathway (which expresses LTD). The antagonism of D1 receptors reduced the amplitude of EPSCs mediated by GluN2B-but not GluN2A-containing NMDARs in the contralateral pathway; this modulatory effect was not observed in the ipsilateral pathway. Our observation that the D1 antagonist equally reduced NMDAR-mediated EPSCs either before or after blockade of GluN2A-and GluN2B-containing NMDARs suggests that, despite the lack of D1 modulation of GluN2B-or GluN2A-containing NMDARs at ipsilateral synapses, other subunits like GluN2C and GluN2D may be subject to D1 modulation (Zhang et al., 2014). In contrast to the lack of D1 modulation, D2 receptors affected the amplitude of GluN2B-containing NMDARs-mediated EPSCs at ipsilateral synapses, but not at contralateral synapses despite the presence of GluN2B-containing NMDARs; indeed, the antagonism of GluN2B-containing NMDARs resulted in a larger reduction of the amplitude of NMDAR-mediated EPSCs at ipsilateral synapses after D2 receptors were blocked. The modulatory effect of D1 and D2 receptors on NMDAR-mediated cortico-striatal synaptic transmission seems to be permissive rather than instructive, because only their antagonists have significant actions whereas their agonists don’t. Our results also indicate that a tonic basal level of D1 receptor activation at contralateral synapses may serve to stimulate GluN2B-containing NMDARs, whereas that of D2 receptors at ipsilateral synapses may inhibit them. Considering the role of GluN2B-containing NMDARs in cortico-striatal plasticity, we predict that D1 receptors will be involved in LTP in the contralateral pathway, while D2 receptors will be involved in LTD in the ipsilateral pathway. Future experiments are needed to further determine how D1 and D2 receptors regulate LTP and LTD through their modulation of GluN2B-containing NMDARs.

Studies in the hippocampus suggest a direct interaction of D1 receptors with GluN2A-containing NMDARs, and D2 receptors with GluN2B-containing NMDARs in a specific complex (Lee et al., 2002; Liu et al., 2006). To achieve the specificity of the modulatory effect we observed at the cortico-striatal synapses, D1 and D2 receptors need to selectively interact with GluN2B-containing NMDARs but not with GluN2A-containing NMDARs. Furthermore, our data indicate that both D1-and D2-expressing MSNs in one pathway contain similar levels of GluN2B-containing NMDARs and are subject to comparable DAergic modulation. How can D1-expressing MSNs that lack D2 receptors be modulated to the same extent as D2-expressing MSNs in the contralateral pathway? The same question can be raised for D2-expressing MSNs in the ipsilateral pathway. Such cross-talk effects have been suggested to result from the extensive recurrent synaptic connections between D1-and D2-expressing MSNs (Gerfen and Surmeier, 2011). In addition to this potential postsynaptic mechanism, different populations of presynaptic cortical neurons could also result in distinct DAergic modulation of MSNs in the ipsilateral or contralateral pathway. Morphological and physiological evidence has revealed two types of cortico-striatal projection neurons: intratelencephalic (IT) neurons that project bilaterally to the striatum, and pyramidal tract (PT) neurons that project ipsilaterally to it (Shepherd, 2013). Future experiments are needed to test if the subunit composition of NMDARs at synapses between IT neurons and striatal MSNs varies depending on the hemispheric origin of the cortical inputs, and whether PT neurons make functionally distinct synapses on ipsilateral MSNs compared to those formed by IT neurons.

Alterations in the ratio of GluN2B-containing NMDARs and GluN2A-containing NMDARs at synapses on MSNs correlate with dysfunctional motor behaviors, which has been suggested to underlie striatum-related neurological disorders such as Parkinson’s disease (PD) and Huntington’s disease (HD) (Gardoni and Bellone, 2015). In an animal model of PD, partial lesions of DAergic fibers had no effect on GluN2B levels, but resulted in an increase of Glu2A levels, while full lesions reduced GluN2B levels without altering GluN2A (Picconi et al., 2004, Gardoni et al., 2006; Paillé et al., 2010). Normalizing the GluN2B/GluN2A ratio with a GluN2A-selective interference peptide, or by pharmacological activation of D1 receptors, restored synaptic plasticity in MSNs and improved motor function (Paillé et al., 2010). In an animal model of HD, a selective enhancement of GluN2B was observed in extrasynaptic NMDARs in striatal MSNs (Zeron et al., 2004; Milnerwood et al., 2010). Intriguingly, overexpression of GluN2B led to increased striatal neurodegeneration (Heng et al., 2009). Considering that altered NMDAR composition underlies striatum-related neurological disorders, our observations of unbalanced GluN2B/GluN2A ratio at ipsilateral *vs.* contralateral cortico-striatal synapses suggest that both pathways may have a distinguishing pathological role in disease etiology and progression.

In summary, we demonstrate that the contralateral cortico-striatal pathway has higher levels of GluN2B-containing NMDARs than the ipsilateral pathway. Such distinct content of GluN2B and GluN2A subunits results in long-term depression in the ipsilateral pathway, and long-term potentiation in the contralateral pathway. NMDAR-mediated synaptic currents in these two pathways are differentially modulated by D1 and D2 receptors. These unexpected findings provide new insights into the mechanisms underlying NMDAR-mediated synaptic transmission at cortico-striatal synapses and have important implications for understanding striatum-related behaviors in healthy and diseased states.

## Acknowledgements

This study was supported by NIH grant 1R21NS-097913 (W.L. and L.P.-M). We thank Karen Ayala-Baylon and Yijian Zhang for mouse colony maintenance, Dr. Mary Phillips for comments on the manuscript, and Dr. Takafumi Inoue (Waseda University, Tokyo, Japan) for data acquisition and analysis software.

## Author Contributions

W.L. and L.P.-M designed experiments and wrote the manuscript. W.L. performed the experiments and analyzed data.

